# Genome Analyses of a New *Mycoplasma* Species From the scorpion *Centruroides vittatus*

**DOI:** 10.1101/394825

**Authors:** Tsunemi Yamashita, Douglas D. Rhoads, Jeff Pummill

## Abstract

Arthropod *Mycoplasma* are little known endosymbionts in insects, primarily known as plant disease vectors. *Mycoplasma* in other arthropods such as arachnids are unknown. We report the first complete *Mycoplasma* genome sequenced, identified, and annotated from a scorpion, *Centruroides vittatus*, and designate it as *Mycoplasma vittatus*. We find the genome is at least a 683,827 bp single circular chromosome with a GC content of 43.7% and with 1,010 protein-coding genes. The putative virulence determinants include 20 genes associated with the virulence operon associated with protein synthesis (SSU ribosomal proteins) and nine genes with fluoroquinolone resistance. Comparative analysis revealed that the *M. vittatus* genome is smaller than other *Mycoplasma* genomes and exhibits a higher GC content. Phylogenetic analysis shows *M. vittatus* as part of the Hominis group of *Mycoplasma*. As arthropod genomes accumulate, further novel *Mycoplasma* genomes may be identified and characterized.

## INTRODUCTION

The investigation of the microbiome has shown important relationships exist between hosts and their symbionts. In eukaryotic genomics projects, symbiont genomes are often revealed via significant variation in GC/AT nucleotide content. In arthropod genomes, Mollicutes as a bacterial class have gained attention due to their reduced genome, symbiotic existence, and their occurrence as pathogens (Thompson et al.2011, Browning and Citti 2014). Within the Mollicutes, the *Mycoplasma* are significant as animal pathogens and arthropod symbionts (Thompson et al. 2011, Leclercq et al. 2014). The current diversity of arthropod *Mycoplasma* is under-represented, but the accumulation of additional arthropod genomes will improve the catalog of *Mycoplasma* diversity as contaminants in arthropod genome assemblies (Leclercq et al. 2014). As Mollicute genomes accumulate, taxonomic revisions with genomic data provide evolutionary insights into these prokaryotes (Hicks et al. 2014, Bolaños et al. 2015). Here, we report a new *Mycoplasma* genome from the striped scorpion, *Centruroides vittatus*, and we designate this *Mycoplasma* as *M. vittatus*.

## MATERIALS AND METHODS

### Genome sequencing and assembly

Total genomic DNA was extracted from three scorpions collected in Pope County, AR with the Qiagen genomic-tip and genomic DNA buffer set (Qiagen, Inc.). The genomic DNA quality was analyzed through 0.9% agarose gel electrophoresis and UV spectroscopy. One genomic DNA sample was sent to the University of Arkansas for Medical Sciences DNA Sequencing Core Facility for library generation and 600 cycle paired-end sequencing on a Illumina MiSeq. Two genomic DNA samples were sent to the National Center for Genome Resources (NCGR, NM) for PacBio 20K library generation and 10 SMRT cell sequencing for each individual genome. The de novo assembly was conducted at the Arkansas High Performance Computing Center at the University of Arkansas. Sequence read data quality control check was using FastQC (0.11.5). The PacBio data was assembled using the Canu Pipeline Assembler (v1.3), while the MiSeq data was assembled using Spades (St. Petersburg genome assembler, ver. 3.6.1). Quality assessment of assemblies was conducted with Quast (Quality Assessment Tool, ver. 4.0). One PacBio assembly (Q1133) was polished with Quiver (PacBio, Inc.,) and also with Pilon (Walker et al. 2014). We aligned our three assemblies against each other with Mauve (V2.4.0) to compare variation among them (Darling et al. 2004). We visualized and compared these draft genomes with CGViewer for blastn comparisons (Grant and Stothard 2008). Prokaryote genome annotation was conducted with the Rapid Annotation using Subsystem Technology (RAST) at http://rasat.nmpdr.org and BASys: www.basys.ca/ (Van Domselaar et al. 2005,Aziz et al. 2008). The genes were categorized into subsystems with the SEED browser <u>at http://www.theseed.org/</u> (Overbeek et al. 2005). Related genomes were identified through RAST and distance matrices generated for selected genomes using the GGDC server for Genome-to-Genome Distance Calculator 2.1 at http://ggdc.dsmz.de/ggdc.php (Meier-Kolthoff et al. 2013). This in-silico reciprocal BLASTn method produces robust distances among bacterial taxa as it incorporates Genome Blast Distance Phylogeny and allows confidence interval estimation. We also included the *M. pulmonis* and *H. crinochetorum* genomes in our RAST and SEED analyses to compare genome and protein features to the *M. vittatus* genome. Phylogenetic relationships were constructed from distance matrices using the Neighbor-Joining algorithm implemented at Trex-online (http://trex.uqam.ca) with the subsequent trees visualized in FigTree v1.4.3 (http://tree.bio.ed.ac.uk/). The genome sequences of the scorpion *Mycoplasma* are available under the GenBank Accession numbers SAMN07809902 and SAMN07809904. Two of the three genomes were identical, thus only unique genomes were submitted.

## RESULTS

### Identification of *M. vittatus* contigs

We assembled three draft *Mycoplasma* genomes from three different scorpions, *Centruroides vittatus,* via data produced as a part of the *C. vittatus* genome project (Yamashita T, Rhoads D, and Pummill J, unpublished data). We identified contigs in our MiSeq and PacBio assemblies from total scorpion DNA that differed markedly in GC content (43.7% GC content vs. 32% in the scorpion genome). There was a single contig in one of the PacBio assemblies (Q1171) of 683,827 bp. The other PacBio assembly (Q1133) contained two contigs of and 511,437 and 163,546 bp (total 674,983). In the MiSeq assembly there were four contigs of 353,646, 177,906, 55,555 and 53,089 bp (total 640,196). The single contig in the Q1171 assembly did not appear to contain any terminal redundancy so it does not appear to represent a complete genome. However, based on Mauve alignments with the two contigs from the Q1133 assembly the Q1171 contig appears to represent a nearly complete assembly (data not shown). Based on the PacBio assemblies the four contigs from the MiSeq assembly were ordered into a single contig with gaps (multiple n residues). Figure 1 shows CGViewer for blastn comparisons and reveals that the two PacBio assemblies differ only for a small region at about 170 bp and reinforces that we have a complete genome between the two assemblies. The MiSeq assembly primarily lacks a region from about 210 to 250 kbp, which is highly conserved between the two PacBio assemblies. This region consists entirely of “hypothetical proteins” according to the RAST annotation and is not recognized as a prophage by the PHASTER server (http://phaster.ca).

**Figure 1.**
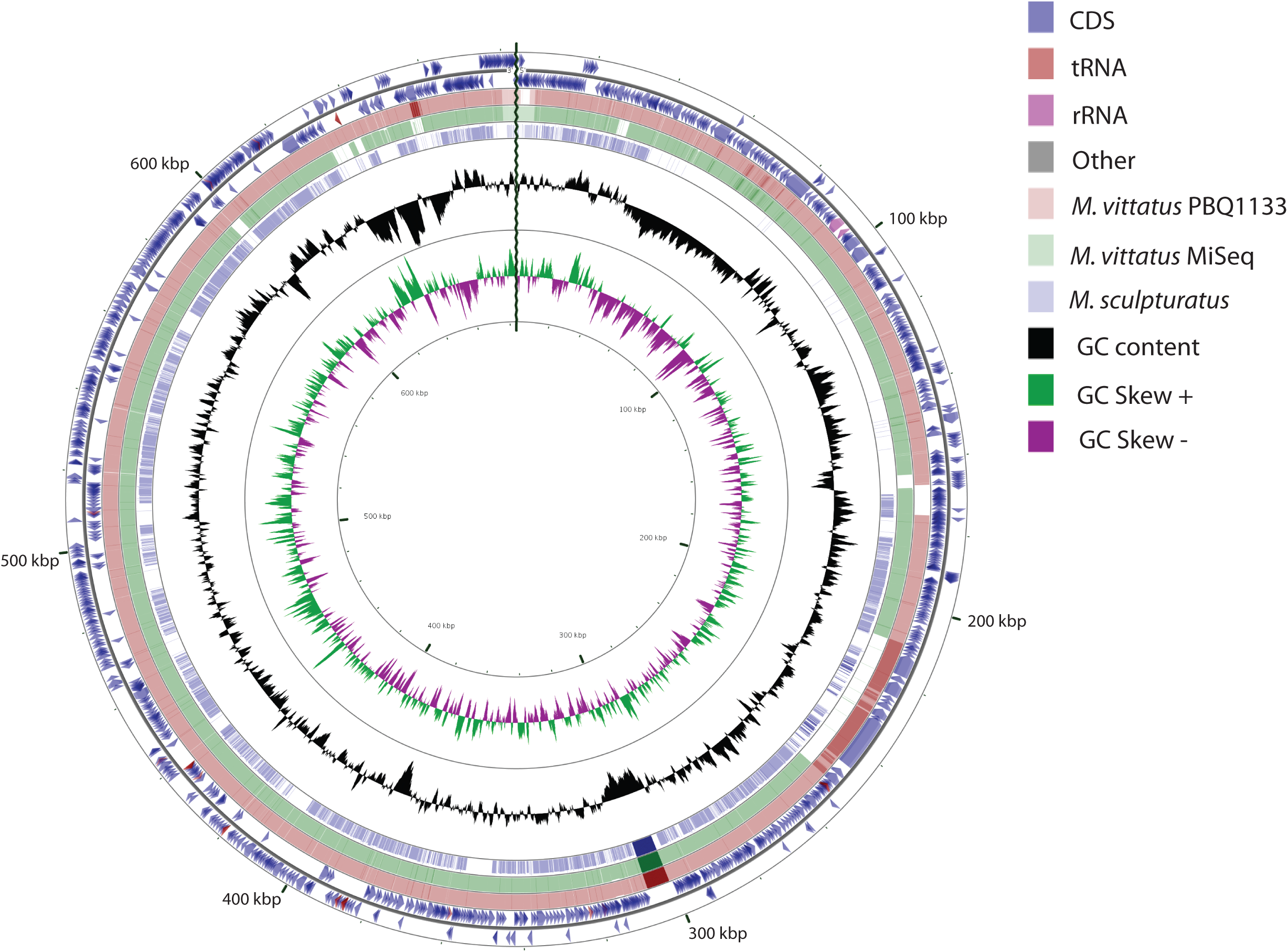
Genome architecture of *M. vittatus* with comparison to *M. sculpturatus*. Moving from outside to inside, the first circle shows the position coordinates for the genome sequence. The second circle shows the predicated locations of the protein coding sequences (in blue) on the plus and minus strands, with tRNA in brown and rRNA in purple. The third pink circle shows the blast alignment of the *M. vittatus* PBQ1133, while the fourth green and fifth blue circles shows the blast alignment of the *M. vittatus* MiSeq and *M. sculpturatus*, respectively. The sixth circle shows the mean centered GC content, with the average GC as baseline and outward projections as higher than average and inwardly projections as lower than average. The seventh ring shows the GC Skew with above zero values in green and below zero values in purple.

### Comparative and evolutionary analysis

We compared the three *M. vittatus* draft genomes to 13 representative mollicute genomes from NCBI. In addition, we extracted 40 contigs representing more than 16 Mbp of likely *Mycoplasma* contigs from a recently released draft genome (NW_019384690.1) for the related scorpion, *Centruroides sculpturatus* (referred to as *M. sculpturatus* and also included in the CGViewer comparison in Figure 1). The Neighbor Joining phylogeny based upon genome distances among 17 genomes based on pairwise distances from the GGDC analyses indicates that *M. vittatus* clusters with the Hominis group of *Mycoplasma* with *M. pulmonis*, the most similar taxon as revealed through our phylogenetic analyses (Figure 2). The comparative genome data are shown in Table S5. The *M. pulmonis* genome is 963,879 bp, 280,052 bp longer than *M. vittatus* and shows 13.9% sequence similarity (1182 identical sites). The RAST server comparison to close strains (i.e., most similar annotated genome) identifies similarities and differences in coding sequences between these two genomes, 434 genes were identified with a known function and 302 genes were identified as similar between the two genomes with 91 genes seen only in *M. pulmonis* and 41 genes in *M. vittatus.*

**Figure 2.**
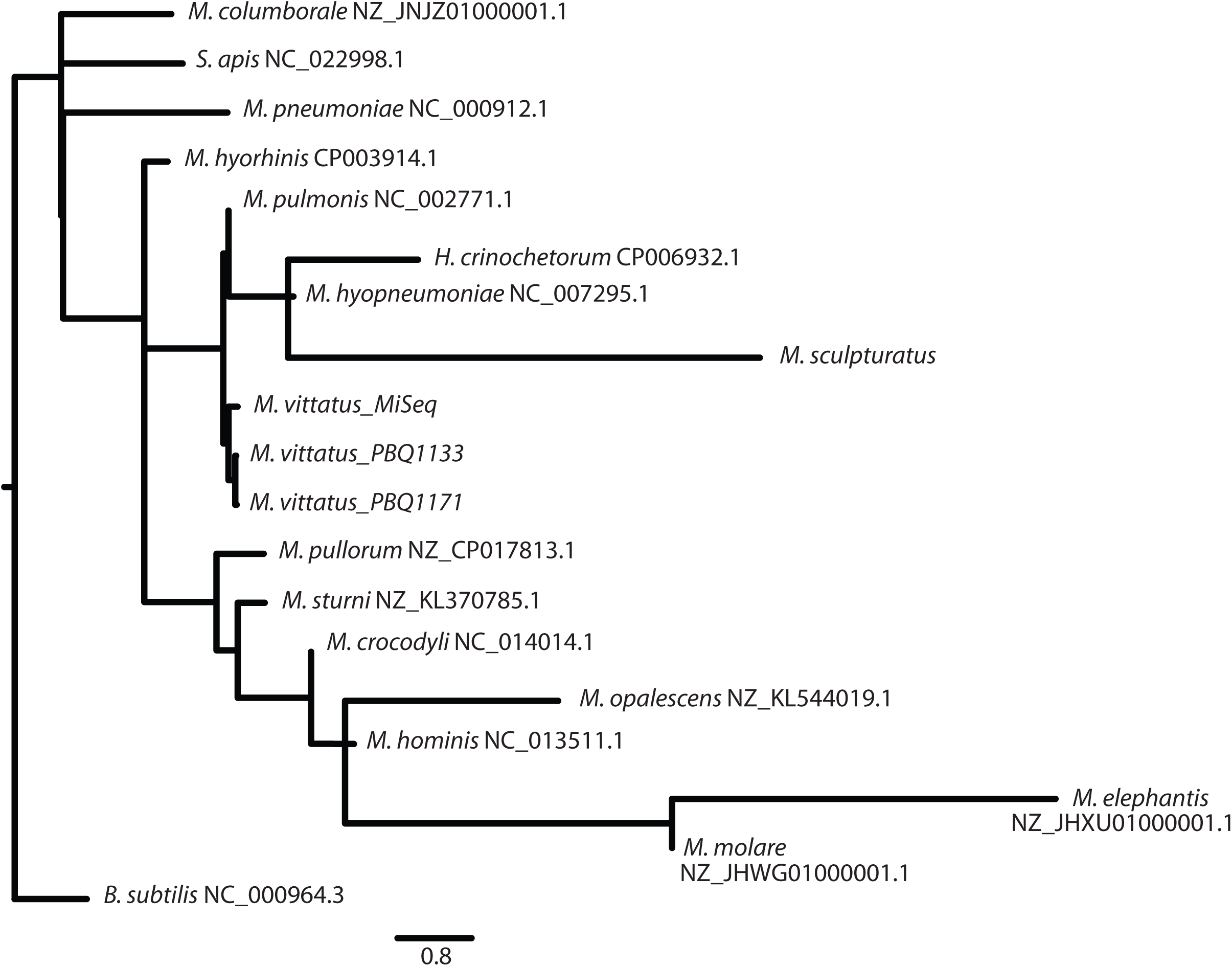
A phylogenetic trees created from GGDC output distances in a Neighbor-Joining tree. The scorpion *Mycoplasma* fall within the hominis group of *Mycoplasma*. The scale represents the number of substitutions per site.

### Genome features

The *M. vittatus* genome from one of the PacBio assemblies contains a 683,827 bp single contig with 43.7% GC content, which appears to deviate from the low GC content seen in other *Mycoplasma* (Figure 1). A total of 1,010 protein coding genes were identified that occupy 88.83% of the genome. Table S1 shows the summary of the genome content in the *M. vittatus*. The non-coding RNAs include 29 tRNA’s one large subunit RNA, and one small subunit RNA (Table S1). For the protein coding genes, 377 (34%) were classified into 18 subsystems and 673 genes (66%) were classified as unknown function (Table S2). Figure S1 shows a comparison of genomic structure among the three *M. vittatus*. Table S5 shows the genome comparison of *M. vittatus* to *M. pulmonis.*

Eleven genes were identified for membrane transport with the bulk associated with ABC transporters (5 genes). Two other systems identified in transport were protein cytoplasmic membrane translocation (4 genes) and two cation transporters (Table S3).

We identified 20 genes associated with virulence, disease, and defense (Table S3). Eleven genes were those associated with the *Mycobacterium* virulence operon and nine genes associated with fluoroquinolone resistance. Ten genes associated with the virulence operon were clustered into three genomic regions: one with three RNA polymerase genes, a second with two LSU ribosomal protein genes with a translation initiation factor, and a third with two SSU ribosomal proteins with two translation factor G genes. Interestingly, no genes associated with adhesion, toxins, nor antibacterial peptides were identified. However, three genes associated with CRISPR proteins were identified: Cas1, Cas2, and Csn1, along with a CRISPR guide cluster.

*Mycoplasma* are known for reduced biosynthetic activities. We identified 298 genes for metabolic activities, but only two genes associated with the respiratory dehydrogenases were identified (Table S4). No genes associated with the ATP synthase nor the electron accepting reactions were identified. The remainder were listed in the following categories: potassium metabolism 6, RNA metabolism 44, protein metabolism 114, DNA metabolism 44, respiration 2, Lipid metabolism 16, carbohydrates metabolism 71, and sulfur metabolism 1.

## DISCUSSION

The *M. vittatus* genome is substantially smaller than *M. pulmonis*, but slightly larger than the genome of *H. crinochetorum* (657,101bp). With a GC content of 42.70%, it also exhibits a higher GC content than other *Mycoplasma* with GC contents between 20% to 40% (Thompson et al. 2011).

As *Mycoplasma* exist primarily as intracellular symbionts (Chen et al. 2017), transporter systems are crucial in obtaining nutrients from their hosts. In the *M. vittatus*, 5 of 11 genes identified in the transporter system fall into the ABC transporter system (Table S3). In *H. crinochetorum* only five genes were identified through the SEED subsystem analysis in the transporter system with none associated with ABC transporters: three genes were identified as involved with protein translocation across cytoplasmic systems and two as cation transporters. *M. pulmonis* exhibited 22 genes associated with membrane transport with the bulk (11/22) in the ABC transporter category.

*M. vittatus* shows 20 genes in virulence, disease and defense (Table S3). Nine genes produce proteins to resist fluroroquinolones, namely in DNA replication (Gyrase and Topoisomerase subunits). The rest are associated with invasion and intracellular resistance. In *H. crinochetorum* 14 genes are virulence, disease, and defense with four genes associated with fluroroquinolone resistance (Gyrase and Topoisomerase subunits), eight in invasion and intracellular resistance, one in Copper homeostasis, and one in multidrug resistance efflux pumps. *M. pulmonis* exhibited 14 genes associated with virulence, disease, and defense with the bulk (9/14) in the invasion and intracellular resistance subsystem.

Our phylogenetic analysis indicates *M. vittatus* is nested in the Hominis group of *Mycoplasma* along with a distinct Mycoplasma in the related scorpion *C. sculpturatus* (Figure 2). This phylogeny mirrors other *Mycoplasma* phylogenies (Thompson et al. 2011, Liu et al. 2012, Hicks et al. 2014). In comparison with the nearest similar species, *M. pulmonis, M.vitattus* exhibits major variation in genome size, gene number, GC content, and average gene length. In fact, *M. vittatus* possesses a genome significantly smaller than those of other species within the Hominis group in our phylogenetic tree (683,827bp vs mean=887,259), and suggests arthropod *Mycoplasma* may house reduced genomes when compared to those symbiotic with vertebrates. These differences also suggest complex evolutionary histories and selection pressures are needed to produce diverse genomes in the *Mycoplasma* phylogeny. Further genomic studies in arthropod genomics should reveal other intricate relationships between arthropods and their *Mycoplasma* symbionts.

## ACKNOWLEDGEMENTS

This publication was made possible by the Arkansas INBRE program, supported by a grant from the National Institute of General Medical Sciences, (NIGMS), P20 GM103429 from the National Institutes of Health.

Author Contributions: Conceived and designed the experiments: T.Y. and D.R. Performed the experiments: T.Y., D.R., and J.P. Analyzed the data: T.Y., D.R., and J.P. Contributed reagents/materials/analysis tools: T.Y., D.R., and J.P. Wrote the paper: T.Y., and D.R. All authors read and approved the final manuscript. The authors declare that they have no competing interests. The funding sponsors had no role in the design of the study; in the collection, analyses, or interpretation of data; in the manuscript writing, and in the publication decision.

